# Acclimation response type shapes temporal stability of cyanobacterial communities under thermal fluctuations

**DOI:** 10.64898/2026.08.07.743510

**Authors:** Arunima Sikder, Pauline Witsel, Frederik De Laender

**Affiliations:** University of Namur; UNamur

**Keywords:** acclimation, community dynamics, cyanobacteria, response diversity, heatwaves, temporal stability

## Abstract

Thermal fluctuations increasingly take the form of recurring heatwaves, yet how the acclimation responses of community members shape collective temporal stability remains untested. Using communities of the marine pico-cyanobacterium *Synechococcus* sp. assembled in microcosms to span a broad range of mean response and response diversity, we tracked total cell density and per-cell chlorophyll *a* through a 16-day warming–cooling fluctuation regime. Temporal stability of total density was predicted by community mean response and response diversity, but only when computed from acute acclimation responses, not when computed from chronic responses at sustained conditions. Higher acute response diversity increased stability, while a higher mean acute response reduced it, independently of intraspecific richness. Community growth was sub-additive at warming transitions but matched the additive prediction at cooling, indicating that inter-strain interactions suppress community growth specifically as the community enters the warm state. Acclimation response is thus a determinant of both the predictability and the realized dynamics of microbial communities under recurring thermal stress.

## Introduction

Climate change is rapidly altering the variance and abruptness of thermal environments [1], making thermal fluctuations a primary ecological driver for biological communities [2, 3]. Thermal variability increasingly takes the form of discrete heatwaves: pulses of elevated temperature that arrive, persist for days, and then recede towards baseline [4, 5]. A common feature of heatwaves is that they recur, so communities experience repeated cycles of warming and cooling rather than a single event [6]. Therefore, a community needs to adjust its performance in two ways within a single heatwave cycle: enduring the elevated temperatures and the transitions into and out of it. A community’s response to such fluctuations is captured by their temporal stability: the constancy of community-level properties such as total biomass or abundance through time [7–9]. Less stable communities may lead to unpredictable biogeochemical processes and have consequences for food webs [10–14]. Marine picocyanobacteria such as *Synechococcus* sp. are major primary producers in oceans worldwide. How they withstand recurring fluctuations thus determines the functioning of the marine ecosystems they underpin.

A key mechanism underlying the temporal stability of biological communities is response diversity, defined as the variation in species’ responses to environmental change. A greater response diversity promotes stability as declines in some species are compensated by positive or neutral responses of others, thereby stabilizing aggregate ecosystem properties (the insurance hypothesis; [9, 15]). To date, numerous theoretical and empirical studies have identified a positive relationship between response diversity and temporal stability [15–17]. Response diversity, however, describes only how species’ responses differ from one another, not their shared direction: when an environmental driver elicits predominantly positive responses across species, a larger mean response synchronizes changes in community biomass, and may reduce stability even when response diversity is high. We therefore expect greater mean responses to lower temporal stability, independent of the stabilizing effect of response diversity, such that the two act jointly rather than separately, a coupling that recent theory shows is regime-specific, differing between pulse-like and press-like disturbance [18]. However, both response diversity and mean response have so far been characterized exclusively as properties of the response elicited by a current pulse of environmental change. This leaves the contribution of past environments unaddressed. Yet response diversity is fundamentally a property of functional traits, and traits are not fixed at the moment of measurement: they can be shaped by conditions an organism experienced beforehand and carried forward as ecological memory or legacies [6, 19, 20]. If response diversity is measured through trait variation, and prior environments can themselves generate trait variation via acclimation, then acclimation history is a candidate source of response diversity that current-pulse measurements cannot capture, with direct but hitherto unknown consequences for temporal stability. Acclimation is widely defined as the reversible adjustment of traits such as growth rate or resource use in response to an environmental condition [21, 22]. The trait shift an organism carries forward from this process is its acclimation response, which can reshape the aggregate properties of a community and thereby its temporal stability under subsequent fluctuations.

An acclimation response can be categorized as either acute or chronic [23]. Both response types are comparative by nature: they compare performance under warm and baseline temperatures. The distinction lies in the environment immediately preceding the measurement. An acute response measures performance immediately after a temperature switch, following acclimation to the opposite temperature. A chronic response measures performance under a sustained temperature, following acclimation to that same temperature. An organism is characterized by both, and the two need not align: an acute response can differ from, or even oppose, a chronic one [24]. Thus, a community can be described by either response type. In a recurring heatwave, this distinction carries important ramifications for stability, because the two responses describe different phases of the cycle: chronic responses reflect differences in physiology during the sustained plateaus between switches, whereas acute responses reflect differences during the transitions into and out of the warm state. Existing knowledge offers no consensus on which matters more for stability.

Inter- and intraspecific diversity are also important determinants of temporal stability [9, 15, 25–27]. The stabilizing effect of richness is thought to arise because richer communities encompass a broader distribution of species responses to environmental change, making richness an indirect proxy for response diversity. However, if response diversity generated by acclimation responses can be measured directly, richness’s classical stabilizing role should, in principle, be absorbed by that direct measure. In such cases, it remains unknown whether richness interacts differently with acute and chronic response types under fluctuating thermal regimes.

To investigate these gaps empirically, we use *Synechococcus* sp. as a biological system with rich intraspecific diversity and variance of thermal preferences [28]. Because *Synechococcus* communities form the productive base of marine ecosystems [29, 30], their stability under thermal fluctuation propagates to marine food webs and biogeochemical cycles. Using microcosm communities that varied in mean response and response diversity computed from acute and chronic acclimation responses, and intraspecific richness, we tracked total cell density and per-cell chlorophyll *a* through a warming–cooling fluctuation regime. Specifically, we ask: (1) does the predictive power of response diversity and mean response for temporal stability depend on acclimation response type (acute or chronic)? (2) does this effect differ between community abundance (total cell density) and a proxy for per-cell physiological state (cellular chlorophyll-*a* content)? (3) does it depend on intraspecific richness? (4) is community stability under fluctuation recoverable from the independent responses of the constituent strains, or does it depart from this additive expectation?

## Materials and methods

We used 12 strains of the marine pico-cyanobacterium *Synechococcus* sp., selected based on intraspecific variation across Atlantic Ocean habitats (Supplementary Table 1) and obtained from the Roscoff Culture Collection, France. Strains were maintained in 50 mL Erlenmeyer flasks of PCR-S11 medium [31] in thermostatic cabinets (Lovibond) on platform shakers (Fisher Scientific) at 150 rpm under a 12h:12h light: dark cycle. The control temperature (C) was set to 19 °C, the mean preferred temperature across the 12 strains, and the warming temperature (T) to 24 °C (C + 5 °C), approximating a heatwave-magnitude thermal anomaly.

### Experimental design

The experiment comprised three sequential phases (Fig. 1): a 10-day acclimation phase, a 48h cross-exposure (response) phase to quantify acclimation responses, and a 16-day community fluctuation phase. Cell density and chlorophyll *a* were measured throughout by flow cytometry.

**Figure 1.**
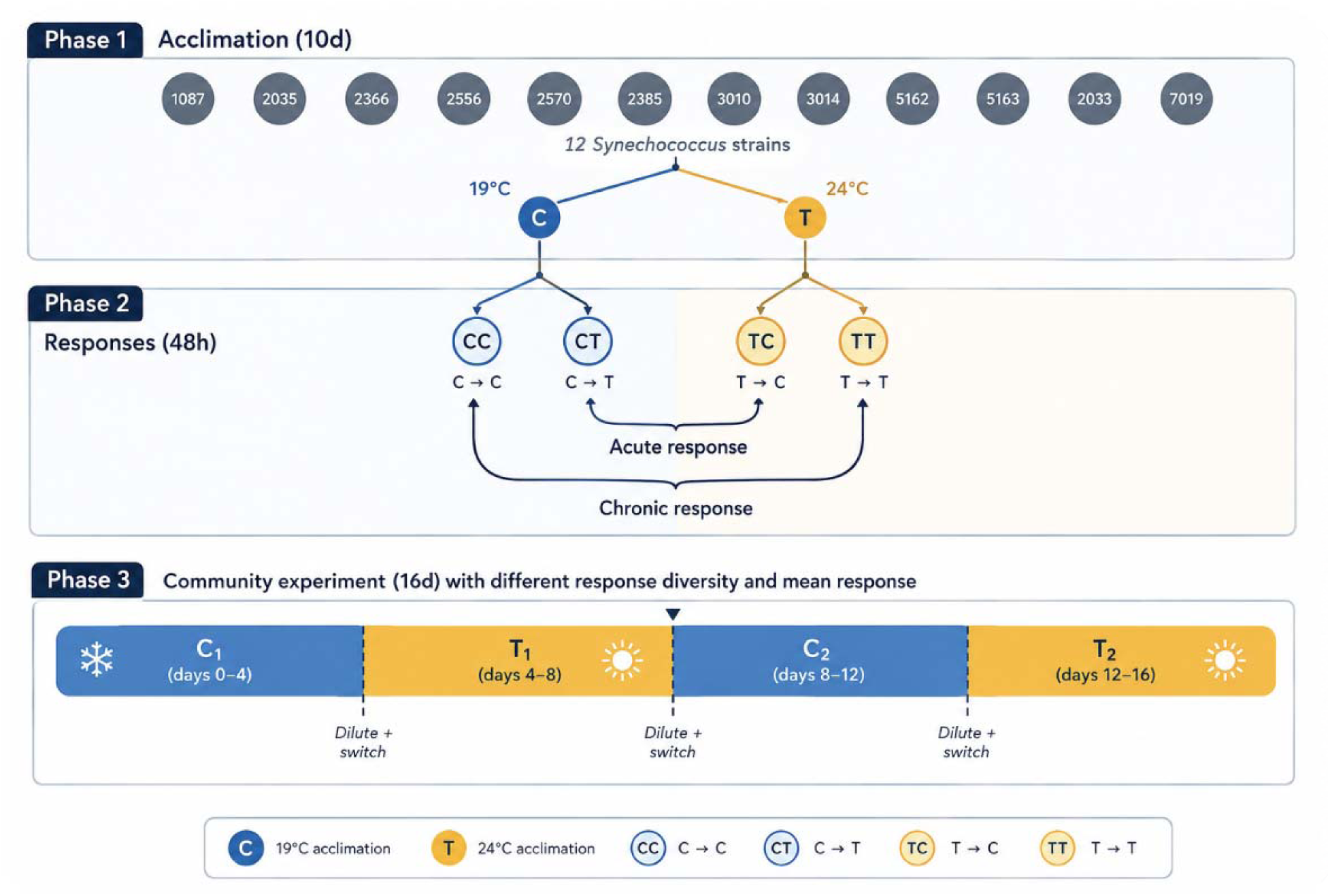
Experimental design. The experiment proceeded through three sequential phases. Phase 1 – Acclimation of 12 strains of *Synechococcus sp*, Phase 2 - Cross-exposure, and Phase 3 – thermal fluctuation. In acclimatio phase, blue circle denotes control temperature (19 °C) and yellow circles denote warming (T, 24 °C). In response phase, the border colors denote the acclimation condition and inner circle color denotes cross-exposure condition.

In phase 1 (acclimation), each of the 12 strains was grown as a monoculture under two thermal regimes (C and T), generating strains with distinct acclimation histories. Monocultures were maintained for 10 days with density measured every 24h (n = 3 replicates; 72 experimental units). In phase 2, we measured the acute and chronic response of each acclimated monoculture (Supplementary Table 2 – Fig. 1). We measured the chronic response by holding the culture at its acclimation temperature. We measured the acute response by switching to the alternate temperature for 48 h, giving four acclimation × cross-exposure combinations (CC, CT, TC, TT; n = 3 per strain × exposure, 144 experimental units). Density was measured every 24 h, and the intrinsic growth rate (*r*) was estimated for each strain in each combination. From these four growth estimates we computed, for each strain, two signed response values. The chronic response was the difference in mean intrinsic growth rate (averaged across replicates) between sustained warming and sustained control, (*r*) TT − CC, and the acute response was the corresponding difference between a warming switch and a cooling switch, (*r*) CT − TC. Each strain was thus characterized by two response values, one for each response type.

#### Community assembly

Communities were assembled to vary in mean response and response diversity, and intraspecific richness (2 or 3 strains). For all possible two- and three-strain combinations, we calculated the mean response and response diversity across constituent strains. We then selected a subset of 40 communities (n = 1) that collectively spanned a broad range of mean response and response diversity values. We added to this selection a monoculture of each strain as a control (n = 3; 36 monoculture units; Supplementary Table 3). All communities were assembled from a common warm-acclimated (C→T) starting state and at equal initial density, so that variation in their subsequent dynamics could be attributed to the response properties of their constituent strains rather than to differences in initial composition or assembly history.

In phase 3 (fluctuation), the resulting 76 experimental units (40 communities and 36 monoculture controls) were exposed to a thermal fluctuation regime over 16 days, structured as four consecutive 4-day blocks alternating between control and warming (Fig. 1): C□ (days 0–4) → T□ (4–8) → C□ (8–12) → T□ (12–16). Cell density and chlorophyll a were measured every 48 h.

#### Sampling and flow cytometry

At each sampling point, a 200 μL aliquot from each well was aseptically transferred to a 96-well plate (Corning Costar) and replaced with 200 μL of pre-equilibrated medium to maintain volume. During Phase 3, all experimental units were diluted by a factor of 3 (25 000 cells μL ¹), every 96 h (days 4, 8, 12), immediately before each environmental switch, with pre-equilibrated medium added to maintain the 6 mL well volume. Cell density and a proxy of per-cell chlorophyll *a* were quantified on a Guava EasyCyte HT-7 cytometer (488 nm excitation; forward scatter for counts, RED-B channel 610/20 nm for chlorophyll). Cell counts were derived from .fcs files after debris removal; sample volume was computed from instrument flow-rate constants. Total density per well (cells mL ¹) was scaled to the 6 mL well volume.

### Statistical analysis

#### Quantifying mean response and response diversity

For each of the 40 communities, mean response and response diversity were computed separately from the acute and chronic response values of their constituent strains (obtained in Phase 2). Mean response was the arithmetic mean of the constituent strains’ signed response values, and response diversity was quantified as the mean pairwise Euclidean distance among constituent strains’ response values [26] :

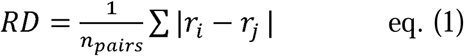

where *r_i_* and *r_j_* are the acute (or chronic) response values of strains *i* and *j*, and the sum is taken over all *n_pairs_* (strain pairs) within a community. This gave each community four composite predictors: mean and response diversity, computed separately from acute and from chronic strain-level responses (Supplementary Table 4), in addition to its intraspecific richness (2 or 3 strains). Predictors were mean-centered prior to modelling.

#### Computing temporal stability

For each community, we quantified temporal stability as the coefficient of variation (CV) of total density and chlorophyll *a* across the 16-day fluctuation regime as

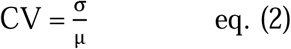

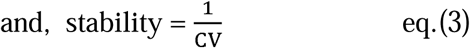

where *σ* is the standard deviation of the response variable and *µ* is its mean.

Because environmental block and time are confounded by design, inference was based on this integrated per-replicate metric rather than on within-regime dynamics. All analyses were performed in R (v4.5.2; R Core Team), with data handling throughout using the *tidyverse* suite (v 2.0.0). Candidate distributions (normal, gamma, lognormal) were compared by ΔAIC for both variables (Supplementary Table 5 – Fig. 2); a lognormal distribution provided the best fit in both cases, and CV was accordingly modelled on the log scale with ordinary least squares. Separate models were fit for predictors derived from the acute and from the chronic response, to test whether acclimation response type determines the predictive value of mean response and response diversity (RD) for temporal stability:

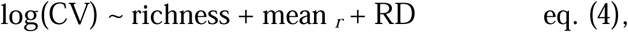

fit independently for density and chlorophyll, and independently using acute-derived and chronic-derived predictors. Richness × predictor interactions were tested in the full model and found non-significant for both response variables (density: p = 0.853; chlorophyll: p = 0.662); richness was therefore retained as an additive covariate only.

**Figure 2.**
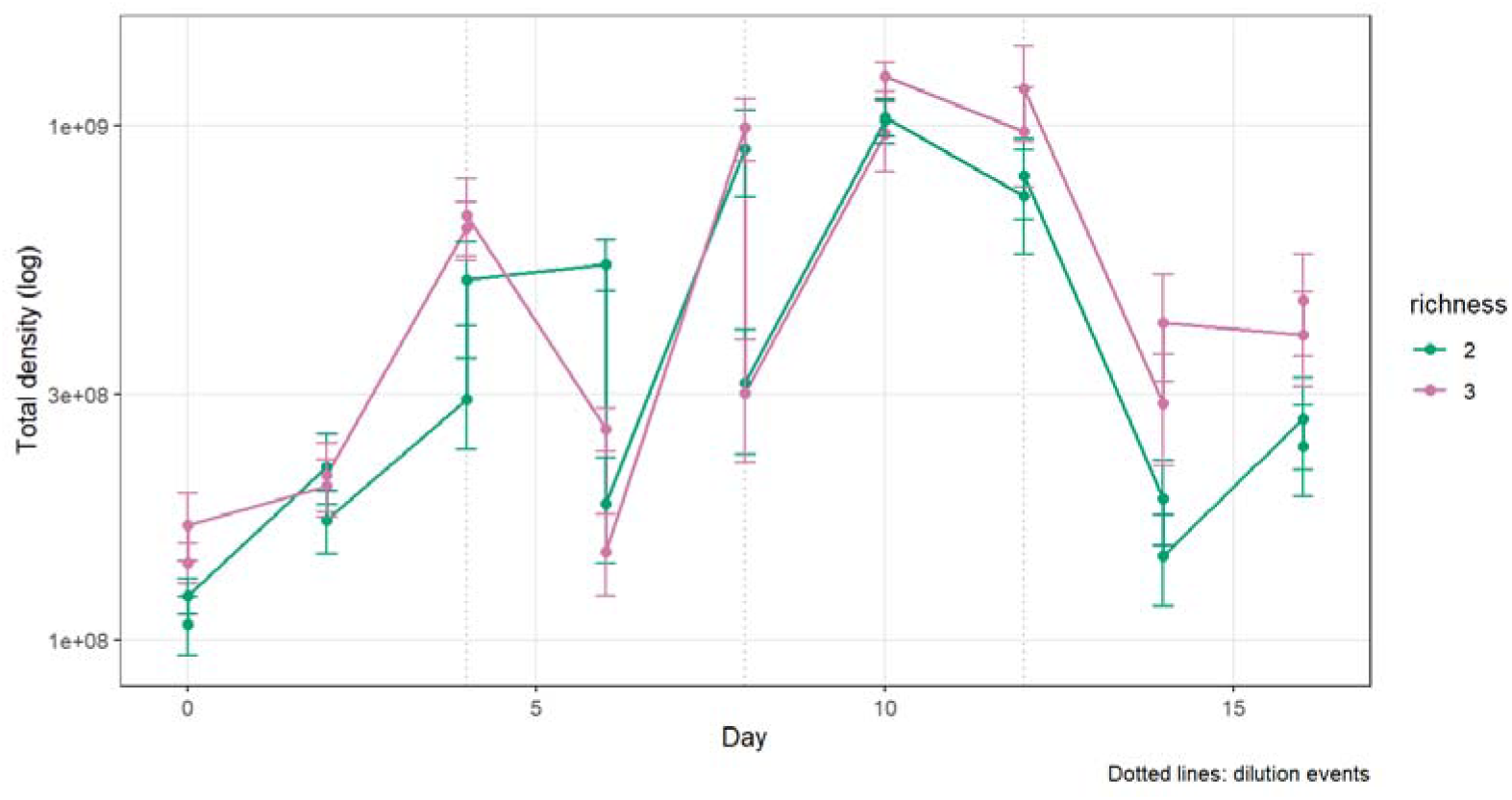
Total community density dynamics and temporal stability throughout the temperature fluctuation regime. Mean total density (log scale) across the 16-day fluctuation regime for communities of two (green) and three (pink) strains. Points show means ± SE averaged across all communities of each richness class at each sampling point; dotted vertical lines mark the three dilution events (approximately 3-fold, on days 4, 8, and 12) immediately preceding each thermal switch. The regime alternated between control (C, days 0–4, 8–12) and warming (T, days 4–8, 12–16) blocks, producing the characteristic sawtooth pattern of growth and dilution-driven decline.

Model diagnostics included Shapiro-Wilk tests on residuals and variance inflation factors (VIF) to confirm collinearity between richness, mean response and response diversity did not compromise coefficient estimates (Supplementary Table 6; all VIF ≤ 1.07). Confidence intervals for all model coefficients were obtained by bias-corrected and accelerated (BCa) bootstrap, resampling communities with replacement over 2000 iterations (*boot* v1.3-32), and are reported alongside coefficient estimates and *p*-values in Figures 3–4.

**Figure 3.**
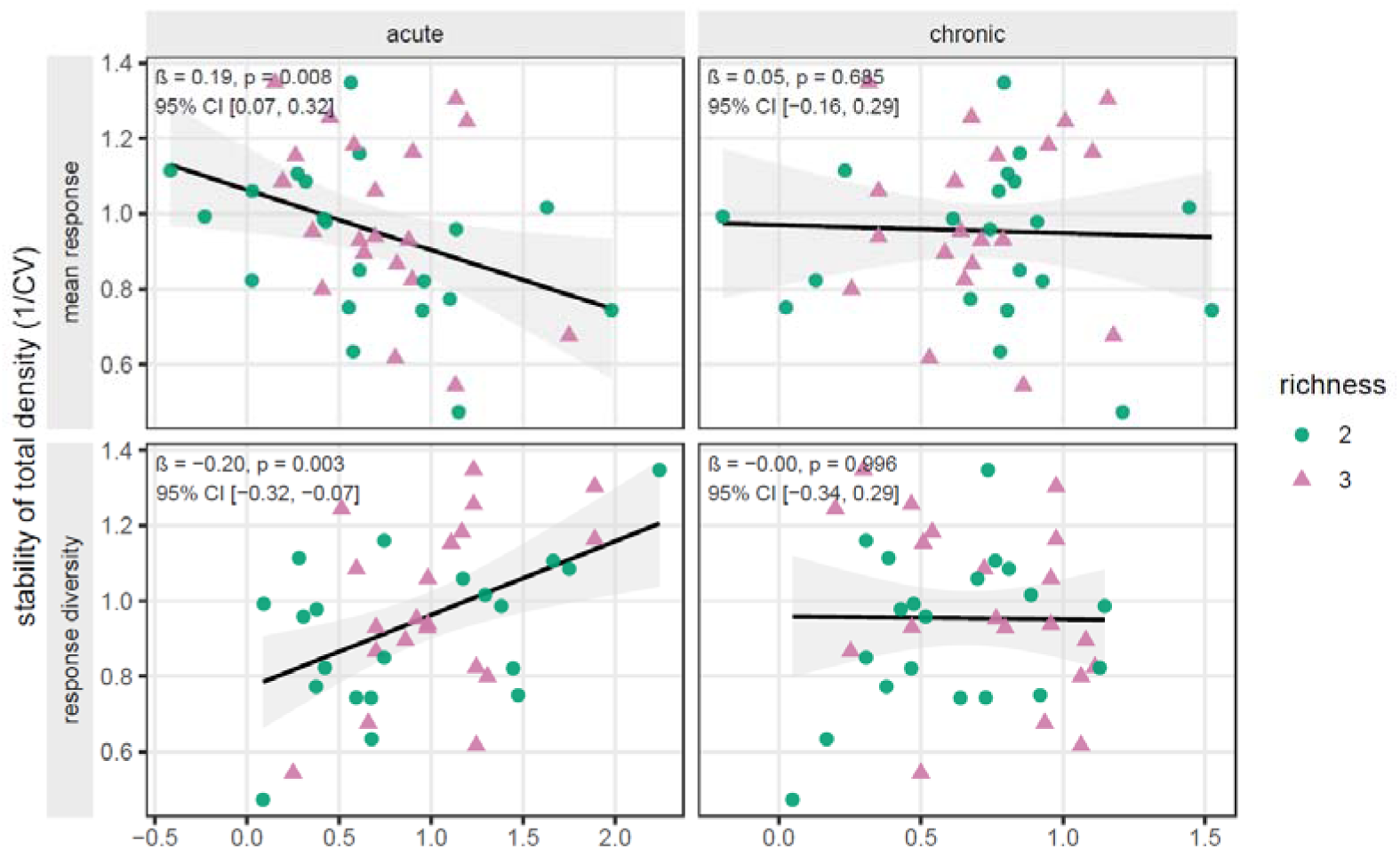
Acute, but not chronic, acclimation responses predict the temporal stability of total density. Temporal stability (reciprocal of the coefficient of variation, 1/CV, of total density across the 16-day fluctuation regime) is plotted against community mean acclimation response (top row) and response diversity (bottom row), computed from acute (left column) or chronic (right column) strain-level responses. Each point represents one of the 40 experimental communities; colour and shape indicate intraspecific richness (green circles, 2 strains; pink triangles, 3 strains). Black lines show ordinary least-squares fits from the model, fit separately for acute- and chronic-derived predictors; shaded bands are 95% confidence intervals on the fitted line. Annotations give the slope (β, on the log-CV scale), p-value, and bias-corrected and accelerated (BCa) bootstrap 95% confidence interval for each predictor.

**Figure 4.**
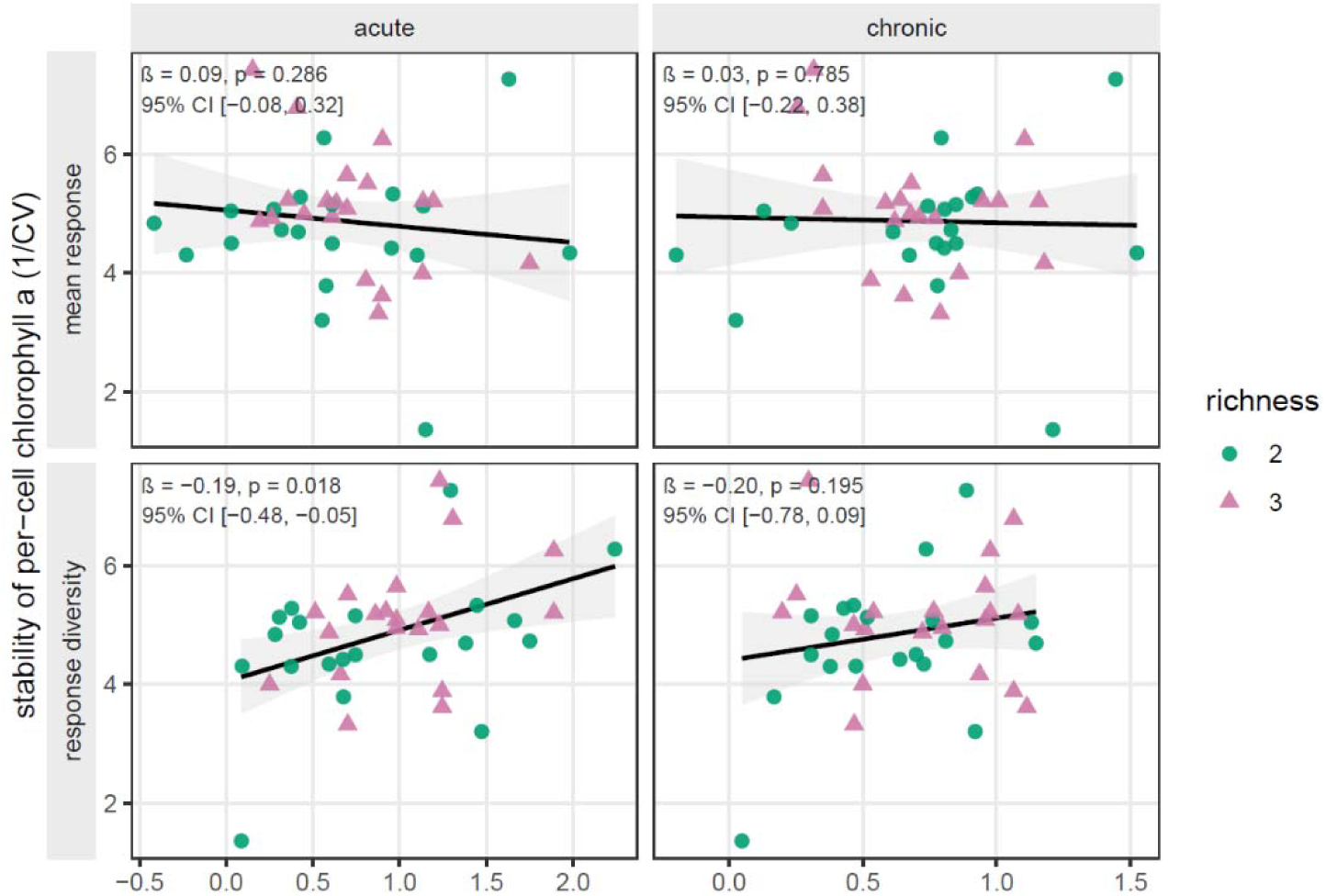
Acclimation responses do not robustly predict the temporal stability of per-cell chlorophyll a. Layout, symbols, colours, and fitted regression lines are as described in Figure 3, with temporal stability expressed as the reciprocal of the CV of per-cell chlorophyll *a* (1/CV). Annotations give β, p, and BCa bootstrap 95% CI for each predictor from the model, fit separately for acute- and chronic-derived predictors.

#### Prediction of temporal stability

To test whether community stability reflected the independent responses of its constituent strains or emerged through their interaction, we compared observed community growth to an additive prediction built from the strains’ monoculture growth rates, which retains each strain’s acclimation response but excludes interaction between strains.

For each community we computed an observed per-capita growth rate (*ρ_o_*) over three transitions: first warming (C→T), cooling (T→C), second warming (C→T):

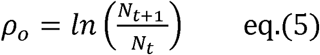

where *N_t_* and *N_t_*_+1_ are community densities at the start and end of the 48 h transition window. Monoculture rates *ρ_p_* were computed over the same windows and combined as

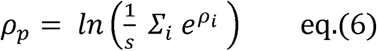

the growth rate of a non-interacting, equal-density mixture of S strains. Rates are averaged as growth factors (in density space) rather than as log rates, since only the former gives the non-interacting expectation (Supplementary Text 1). Within each transition, we regressed observed community growth (*ρ_o_*) on the additive prediction (*ρ_p_*) across all communities; departure of the fitted slope from unity or the intercept from zero indicates that the additive prediction is biased for that transition. To quantify the direction and size of that departure, we computed the signed departure of each community from its prediction (*ρ_o_* – *ρ_p_*) and tested, separately for each transition, whether its mean differed from zero using a linear mixed model with a random intercept nested for community and replicate. A negative mean departure indicates that communities grew more slowly than a non-interacting mixture of their strains would predict (sub-additivity), a positive departure that they grew faster.

## Results

### High acute response diversity and low acute mean response convey greater temporal stability of total density

Temperature fluctuations drove large changes in total community density across the regime (Fig. 2). Across communities, the temporal stability of total density was predicted by the mean response and response diversity of their constituent strains, but only when these were computed from acute acclimation responses.

On the log-CV scale, CV increased with acute mean response (β = 0.186, p = 0.008) and decreased with acute response diversity (β = −0.200, p = 0.003; Supplementary Table 7): communities were thus less stable the larger their mean acute response, and more stable the greater their acute response diversity, together explaining a substantial fraction of the among-community variation in stability (R^2^ = 0.36; F□,□□ = 6.61, p = 0.001; Supplementary Table 7).

The same metrics computed from chronic responses did not predict stability (mean chronic p = 0.685; chronic response diversity p = 0.996). Acute mean response and acute response diversity were uncorrelated across communities (r = −0.04), so their opposing effects on stability are separable rather than two expressions of a single axis.

### Temporal stability is unaffected by intraspecific richness

Intraspecific richness did not predict the temporal stability of total density and did not modify the effects of mean response or response diversity: richness × predictor interactions were non-significant (p = 0.853; Fig. 3), and richness was retained as an additive covariate with no independent effect (β = −0.059, p = 0.375; Supplementary Table 7). Because richness is thought to stabilize communities largely by broadening the distribution of responses they contain, its lack of an independent effect here indicates that this stabilizing information is already captured directly by response diversity, leaving no residual contribution once response diversity is measured.

### The temporal stability of per-cell chlorophyll a is not robustly related to acclimation responses

In contrast to total density, the temporal stability of per-cell chlorophyll *a* was only weakly related to acclimation responses (Fig. 4; Supplementary Table 8). Acute mean response did not predict chlorophyll CV (β = 0.089, p = 0.286; Fig. 4). Acute response diversity was associated with lower chlorophyll CV in the full dataset (β = −0.193, p = 0.018), but this rested on a single highly influential replicate (community C2, replicate 3; Cook’s D = 0.67): with that replicate excluded, the association was no longer significant (p = 0.071), and we therefore do not interpret it as robust (both fits reported in Supplementary Table 8). The chronic metrics did not predict chlorophyll CV (mean chronic p = 0.785; chronic response diversity p = 0.195; Fig. 4).

### Community growth is sub-additive at warming transitions

Community growth departed from the prediction of a non-interacting mixture of its constituent strains and did so specifically at the warming transitions (Fig. 5; Supplementary Table 9). At both the first and second warming steps, communities grew substantially slower than predicted (mean departure = −1.26, p = 0.001; and −1.12, p < 10□□), whereas at the intervening cooling step observed growth was closer to the additive prediction (mean departure = −0.17, p = 0.66; Fig. 5). Because the prediction retains each strain’s own acclimation response but excludes any effect of strains growing together, this warming-specific shortfall indicates that inter-strain interactions suppress community growth precisely when the community is driven into the warm state, and not when it relaxes back toward baseline. Non-additivity in these communities is therefore concentrated at the thermal transitions that destabilise them, consistent with the interactions, rather than the strains’ independent responses, shaping community dynamics under warming.

**Figure 5.**
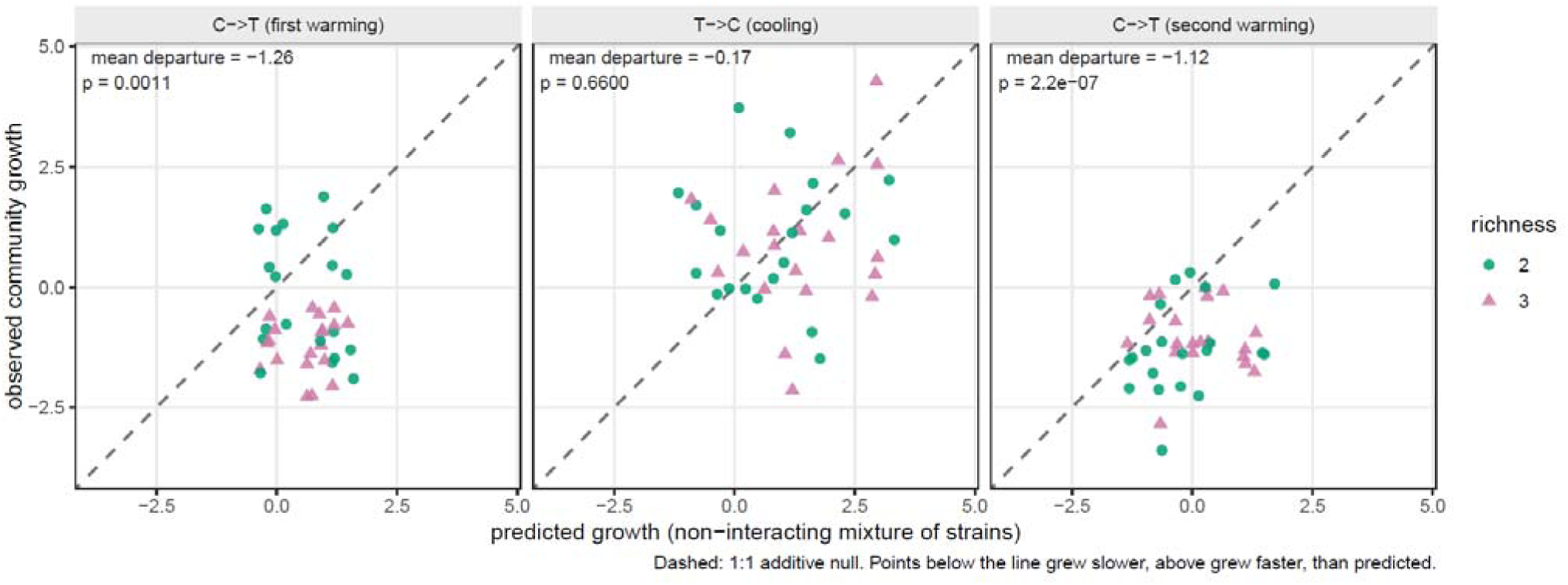
Community growth is sub-additive at warming transitions but matches the additive prediction at cooling. For each of the 40 experimental communities, observed per-capita growth rate (*ρ_o_*) is plotted against the growth rate predicted for a non-interacting, equal-density mixture of the community’s constituent strains (*ρ_p_*), separately for three transitions of the fluctuation regime: first warming (C→T, days 4–6), cooling (T→C, days 8–10), and second warming (C→T, days 12–14). Each point is one community replicate; colour and shape indicate intraspecific richness (green circles, 2 strains; pink triangles, 3 strains). The dashed line is the 1:1 additive null: points falling below it grew more slowly than the non-interacting prediction, points above grew faster. Annotations give the mean signed departure (*ρ_o_* − *ρ_p_*, averaged across all communities in that transition) and its p-value from a linear mixed model testing departure against zero (random intercept nested for community and replicate).

## Discussion

We asked whether the mean response and response diversity resulting from two different acclimation responses predict aggregate temporal stability under a fluctuating thermal regime, and whether community stability is recoverable from the independent responses of its constituent strains. We find that temporal stability of total density is predicted by mean response and response diversity, but only when these are computed from acute acclimation responses, not chronic ones and that departure from the additive expectation is governed instead by which strains are combined, independently of the averaged metrics. Acclimation response is thus a within-population, reversible axis of community variation that has not previously been isolated in microbial community-stability experiments, and it operates on stability at two distinct levels: as an averaged predictor and as an emergent, composition-dependent effect.

### Acute acclimation responses predicted temporal stability

Mean response and response diversity predicted stability when computed from acute but not chronic responses (Fig. 3; Supplementary Table 7), identifying the response expressed during a thermal transition as the axis of acclimation that carries information about stability under a fluctuating regime. This is consistent with the structure of the regime itself: in a sequence of plateaus and switches, the destabilizing events are the transitions, and it is the acute response that quantifies performance across them, whereas the chronic response reflects a physiological state resolved during the intervening plateaus. That greater acute response diversity raised stability while a larger acute mean response lowered it (Fig. 3) matches the two mechanisms directly: divergent responses buffer the aggregate through compensation, whereas a large, shared response synchronizes it [15, 18].

One explanation for the primacy of the acute axis is a trade-off between the magnitude and the rate of plastic adjustment. Plasticity rates vary across taxa, and larger-magnitude acclimation responses are often slower [21, 32]. A chronic response optimizes trait adjustment to sustained conditions but may reduce the speed and reversibility of subsequent adjustment; precisely the properties that matter when transitions are abrupt and repeated. Prolonged acclimation can also increase the variance of physiological rates without improving their mean [33]. We note that this trade-off is a hypothesis: we did not measure plasticity rates directly and testing it would require resolving each strain’s rate and magnitude of adjustment. More broadly, our result extends the beneficial acclimation debate [34, 35]: under a recurring regime, the advantage of the acute over the chronic response may lie in its match to a regime dominated by the transitions [3].

### Richness contributed no stability beyond response diversity

Intraspecific richness did not predict temporal stability, nor did it modify the effect of mean response or response diversity (Supplementary table 7). Because richness is thought to stabilize communities largely by broadening the distribution of responses they contain, its lack of an independent effect once response diversity is measured directly indicates that its classical stabilizing role is in this case completely captured by response diversity. Richness stabilizes through the response diversity it samples; when that diversity is quantified from acclimation responses rather than inferred from strain number, richness did not matter. This is consistent with the broader view that the form of intraspecific variation, rather than its amount, governs stability [36, 37].

### Growth and physiology decoupled under fluctuation

The effect of mean response and response diversity on stability was clear for total density but absent for per-cell chlorophyll *a* (Fig. 4; Supplementary table 8). This decoupling reflects that the two are governed on different timescales: total density integrates the growth response of all strains across every transition, whereas per-cell chlorophyll *a* is a physiological trait regulated rapidly and largely independently of growth over the 48 h intervals. Because growth is determined by many processes of which pigment content is only one, the absence of a chlorophyll *a* effect is consistent with, rather than contradictory to, the effect on density; different aggregate properties of a community need not co-vary under perturbation [7, 8, 38].

Cellular chlorophyll *a* is adjusted in response to temperature and light on timescales of hours to days [39], and in *Synechococcus* sp. the thermal optimum for chlorophyll-based growth differs from that for carbon-based growth [40], so cell number and per-cell chlorophyll *a* are not interchangeable readouts of community response. The acclimation-response effect therefore propagates through the trait that tracks community-level growth dynamics, not through the per-cell physiological state, which both community types regulate similarly.

### Deviation from the additive prediction was greatest during transition to warming

Community growth fell below the non-interacting expectation at both warming transitions but matched it at cooling (Fig. 5; Supplementary table 9) but communities varied widely around the prediction at cooling. Because the prediction retains each strain’s own acclimation response but excludes any effect of strain interactions, this warming-specific shortfall suggests stronger inter-strain interactions during transitions to warming and weaker upon cooling. For *Synechococcus* sp. assemblages under recurring heatwaves, these warming transitions are the phases that accumulate as marine heatwaves intensify [1, 4], so the phases in which community productivity is least recoverable from strain-level responses are also those that a warming ocean will impose most often. This complements the stability result: the acute response metrics predict stability (Fig 3; Supplementary table 7), while the community’s realized growth under warming additionally reflects interactions that strain-level responses do not capture [26, 41].

### Synthesis

Temporal stability under a fluctuating thermal regime depends on the acclimation responses a community’s strains carry, irrespective of intraspecific richness. That dependence operates at two levels: the acute responses of the strains predict stability as a community-level property, while the timing in the cooling-warming cycle determines how predictable growth is for specific strain combinations. For marine pico-cyanobacteria, whose populations carry substantial intraspecific variation, this implies that the stability of the primary production they sustain under recurrent heatwaves depends on acclimation responses. Anticipating how microbial primary-producer communities respond to an increasingly variable climate may therefore require tracking their acclimation histories, not only their composition. Future work should extend these results to longer and more varied fluctuation regimes and to direct measurement of strain interactions.

## Notes

### Competing Interest Statement

The authors have declared no competing interest.

