## Supplementary Materials for "Acclimation response type shapes temporal stability of cyanobacterial communities under thermal fluctuations"

**Table 1. Strain identities and collection metadata: .** Source strains of *Synechococcus* sp. used in this study, obtained from the Roscoff Culture Collection (RCC), Roscoff, France. RCC number gives the accession number of each strain in the collection. Pigment indicates the phycobiliprotein-based chromatic type reported by the collection (3a–3d); a dash indicates no type was recorded, and "3b or 3d" indicates an ambiguous classification in the collection record. Light and Temperature give the culturing conditions reported for each strain, in µmol photons m⁻² s⁻ and °C respectively. Collected from gives each strain's geographic origin as recorded by the collection.

| **RCC number** | **Strain ID** | **Habitat** | **Pigment** | **Light** | **Temperature** | **Collected from** |
| --- | --- | --- | --- | --- | --- | --- |
| 3010 | MICROVIR 16CR_2_clonal | North Sea | 3a | 35 | 13 | UK |
| 3014 | MICROVIR 10CR_4-4_clonal | North Sea |  | 35 | 13 | Norway |
| 1087 | A15-24 | Atlantic Ocean | 3c | 20 | 22 | North Atlantic |
| 2366 | WH8103Syn Clonal | Atlantic Ocean | 3b | 80 | 22 | International waters |
| 2033 | WH8109 | Sargasso Sea | 3b | 20 | 22 | USA |
| 2035 | Syn 20 | Atlantic Ocean | 3a | 80 | 22 | Raunefjorden, Norway |
| 2556 | A15-28 Clonal | Atlantic Ocean | 3c | 20 | 22 | North Atlantic |
| 2570 | MICROVIR 16CH_1-clonal | Atlantic Ocean | 3d | 20 | 22 | UK |
| 7019 | Bergen-MES7-4 | North Sea | _ | 35 | 13 | Raunefjorden, Norway |
| 5162 | MV1715 | Atlantic Ocean | 3a | 20 | 22 | USA |
| 5163 | MV0610 | Atlantic Ocean | 3b or 3d | 20 | 22 | USA |
| 2385 | MICROVIR 10CR_4-3 Clonal | Atlantic Ocean | 3a | 33 | 13 | Norway |

**Table** [1]**2. Strain-level acute and chronic response values.**

Intrinsic growth rate (r) was estimated for each of the 12 Synechococcus strains under four acclimation × cross-exposure combinations (CC, CT, TC, TT). The acute response is CT − TC and the chronic response is TT − CC, each averaged across three replicates. Values are in units of d⁻¹.

| **Strain** | **Acute response (CT − TC)** | **Chronic response (TT − CC)** |
| --- | --- | --- |
| 1087 | 0.9140 | 0.8630 |
| 2033 | 0.6160 | 1.1200 |
| 2035 | 1.2900 | 0.4850 |
| 2366 | 1.1900 | 1.2300 |
| 2385 | 0.9830 | 1.0000 |
| 2556 | 1.1100 | 1.1900 |
| 2570 | −0.1840 | −0.4350 |
| 3010 | 0.2380 | 0.6950 |
| 3014 | 1.6800 | 1.1600 |
| 5162 | −0.5570 | 0.4240 |
| 5163 | −0.2750 | 0.0391 |
| 7019 | 2.2800 | 1.8900 |

**Table 3. Community composition.**

The 40 experimental communities, their constituent strains, and the response axis (acute or chronic) on which they were assembled. Community labels encode the assembly axis (A, acute; C, chronic), and intraspecific richness (2 or 3 strains). Each community label appears five times, corresponding to five distinct strain combinations sharing that label.

| **Richness** | **Replicate** | **Strain composition** | **Assembly basis** |
| --- | --- | --- | --- |
| 2 | 1 | 2035+2570 | acute |
| 2 | 2 | 3014+7019 | acute |
| 2 | 3 | 2570+5163 | acute |
| 2 | 4 | 2385+3010 | acute |
| 2 | 5 | 2385+7019 | acute |
| 3 | 1 | 1087+2385+3014 | acute |
| 3 | 2 | 2035+2385+2570 | acute |
| 3 | 3 | 2366+2570+3014 | acute |
| 3 | 4 | 2385+5162+7019 | acute |
| 3 | 5 | 2570+3014+5163 | acute |
| 2 | 1 | 5162+5163 | acute |
| 2 | 2 | 2556+5162 | acute |
| 2 | 3 | 2033+3010 | acute |
| 2 | 4 | 1087+2035 | acute |
| 2 | 5 | 2366+5162 | acute |
| 3 | 1 | 2033+3010+5163 | acute |
| 3 | 2 | 2035+5163+5162 | acute |
| 3 | 3 | 2556+3010+5162 | acute |
| 3 | 4 | 1087+2366+5163 | acute |
| 3 | 5 | 1087+2035+3010 | acute |
| 2 | 1 | 2385+3010 | chronic |
| 2 | 2 | 3014+5162 | chronic |
| 2 | 3 | 2570+3010 | chronic |
| 2 | 4 | 3010+3014 | chronic |
| 2 | 5 | 2035+2385 | chronic |
| 3 | 1 | 2035+3014+7019 | chronic |
| 3 | 2 | 2556+2385+2570 | chronic |
| 3 | 3 | 3014+5162+7019 | chronic |
| 3 | 4 | 1087+2570+3014 | chronic |
| 3 | 5 | 2035+2385+2570 | chronic |
| 2 | 1 | 2033+2035 | chronic |
| 2 | 2 | 2556+5163 | chronic |
| 2 | 3 | 2366+2556 | chronic |
| 2 | 4 | 1087+3010 | chronic |
| 2 | 5 | 2033+5162 | chronic |
| 3 | 1 | 2033+2035+5162 | chronic |
| 3 | 2 | 2366+2556+5162 | chronic |
| 3 | 3 | 2556+3010+5163 | chronic |
| 3 | 4 | 1087+2035+2366 | chronic |
| 3 | 5 | 2035+2556+3010 | chronic |

**Table 4. Per-community mean response and response diversity.**

Mean response and response diversity (mean pairwise Euclidean distance; Ross et al. 2023) computed separately from acute (CT − TC) and chronic (TT − CC) strain-level responses for each of the 40 assembled communities. These are the predictors used in the stability models (Figs 3–4). All values are centred prior to modelling; uncentred values are shown here. A = acute-assembled; C = chronic-assembled; richness = 2 or 3 strains.

| **Repl** | **Richness** | **Mean acute** | **RD acute** | **Mean chronic** | **RD chronic** |
| --- | --- | --- | --- | --- | --- |
| 1 | 2 | 0.553 | 1.474 | 0.025 | 0.920 |
| 2 | 2 | 1.980 | 0.600 | 1.525 | 0.730 |
| 3 | 2 | −0.230 | 0.091 | −0.198 | 0.474 |
| 4 | 2 | 0.611 | 0.745 | 0.847 | 0.305 |
| 5 | 2 | 1.631 | 1.297 | 1.445 | 0.890 |
| 1 | 3 | 1.192 | 0.511 | 1.008 | 0.198 |
| 2 | 3 | 0.696 | 0.983 | 0.350 | 0.957 |
| 3 | 3 | 0.895 | 1.243 | 0.652 | 1.110 |
| 4 | 3 | 0.902 | 1.891 | 1.105 | 0.977 |
| 5 | 3 | 0.407 | 1.303 | 0.255 | 1.063 |
| 1 | 2 | −0.416 | 0.282 | 0.232 | 0.385 |
| 2 | 2 | 0.277 | 1.667 | 0.807 | 0.766 |
| 3 | 2 | 0.427 | 0.378 | 0.907 | 0.425 |
| 4 | 2 | 1.102 | 0.376 | 0.674 | 0.378 |
| 5 | 2 | 0.316 | 1.747 | 0.827 | 0.806 |
| 1 | 3 | 0.193 | 0.594 | 0.618 | 0.721 |
| 2 | 3 | 0.153 | 1.231 | 0.316 | 0.297 |
| 3 | 3 | 0.264 | 1.111 | 0.770 | 0.511 |
| 4 | 3 | 0.610 | 0.977 | 0.711 | 0.794 |
| 5 | 3 | 0.814 | 0.701 | 0.681 | 0.252 |
| 1 | 2 | 0.611 | 0.745 | 0.847 | 0.305 |
| 2 | 2 | 0.561 | 2.237 | 0.792 | 0.736 |
| 3 | 2 | 0.027 | 0.422 | 0.130 | 1.130 |
| 4 | 2 | 0.959 | 1.442 | 0.927 | 0.465 |
| 5 | 2 | 1.137 | 0.307 | 0.742 | 0.515 |
| 1 | 3 | 1.750 | 0.660 | 1.178 | 0.937 |
| 2 | 3 | 0.636 | 0.863 | 0.585 | 1.083 |
| 3 | 3 | 1.134 | 1.891 | 1.158 | 0.977 |
| 4 | 3 | 0.803 | 1.243 | 0.529 | 1.063 |
| 5 | 3 | 0.696 | 0.983 | 0.350 | 0.957 |
| 1 | 2 | 0.953 | 0.674 | 0.802 | 0.635 |
| 2 | 2 | 0.418 | 1.385 | 0.615 | 1.151 |
| 3 | 2 | 1.150 | 0.080 | 1.210 | 0.040 |
| 4 | 2 | 0.576 | 0.676 | 0.779 | 0.168 |
| 5 | 2 | 0.029 | 1.173 | 0.772 | 0.696 |
| 1 | 3 | 0.450 | 1.231 | 0.676 | 0.464 |
| 2 | 3 | 0.581 | 1.165 | 0.948 | 0.537 |
| 3 | 3 | 0.358 | 0.923 | 0.641 | 0.767 |
| 4 | 3 | 1.131 | 0.251 | 0.859 | 0.497 |
| 5 | 3 | 0.879 | 0.701 | 0.790 | 0.470 |

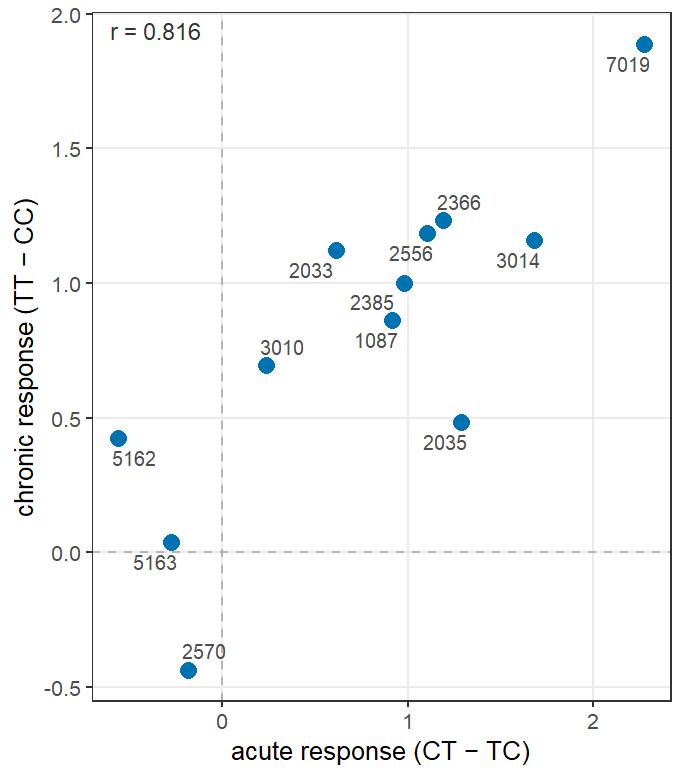

**Figure 1.** Relationship between the acute (CT − TC) and chronic (TT − CC) acclimation responses of the 12 Synechococcus strains used in this study. Each point is one strain; strain codes are labelled. Dashed lines mark zero on each axis. The Pearson correlation coefficient (r = 0.816) indicates that strains with large acute responses also tend to have large chronic responses; both axes share a dominant current-condition sensitivity component. The positive correlation means that the two response contrasts are not independent, and that acute and chronic community-level predictors differ in which biological axis they foreground rather than reflecting entirely separate strain properties.

**Table 5. Candidate distribution comparison for stability (CV) variables.**

Goodness-of-fit statistics and information criteria for four candidate distributions fitted to the coefficient of variation (CV) of total density (n = 40) and per-cell chlorophyll a (n = 40). Lognormal provided the best fit for both variables by AIC and Anderson-Darling statistic and was therefore used as the modelling distribution (ordinary least-squares on log-transformed CV). KS = Kolmogorov-Smirnov; AD = Anderson-Darling. Selected distribution in italics.

| **Variable** | **Distribution** | **AIC** | **BIC** | **KS statistic** | **AD statistic** |
| --- | --- | --- | --- | --- | --- |
| Density CV | Normal | 20.15 | 23.53 | 0.146 | 1.132 |
|  | Gamma | 12.75 | 16.12 | 0.117 | 0.587 |
|  | *Lognormal* | 10.15 | 13.52 | 0.101 | 0.391 |
|  | Weibull | 23.60 | 26.98 | 0.146 | 1.457 |
| Chlorophyll CV | Normal | −74.83 | −71.45 | 0.251 | 4.955 |
|  | Gamma | −101.59 | −98.21 | 0.196 | 2.711 |
|  | *Lognormal* | −111.77 | −108.39 | 0.162 | 1.859 |
|  | Weibull | −78.55 | −75.17 | 0.240 | 5.302 |

**Figure 2. Distribution fit comparison for CV of total density and per-cell chlorophyll a.**

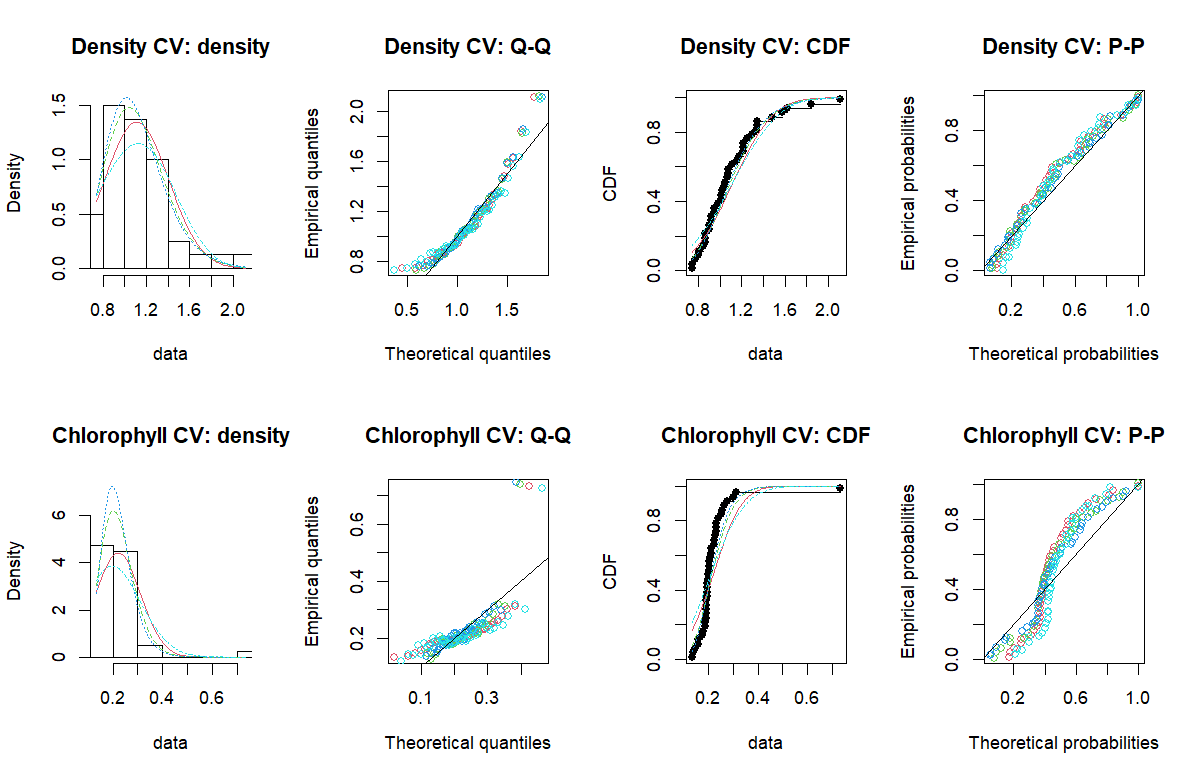

**Figure S2.** Goodness-of-fit comparison for four candidate distributions (normal, gamma, lognormal, Weibull) fitted to the coefficient of variation (CV) of total density (top row) and per-cell chlorophyll a (bottom row), using density plots, Q-Q plots, empirical CDF plots, and P-P plots. The lognormal distribution (cyan) provided the best fit for both variables by AIC and Anderson-Darling statistic (Table S5) and was selected as the modelling distribution, equivalent to applying ordinary least-squares regression to log-transformed CV.

**Table S6. Model diagnostics for the main stability models.**

Shapiro-Wilk test on model residuals, maximum variance inflation factor (VIF) among predictors, and number of observations with Cook's distance > 0.5 for the total-density and per-cell chlorophyll a stability models (log(CV) ~ richness + mean response + response diversity). Chlorophyll residuals departed from normality in the full model (W = 0.864, p < 0.001) due to the influential point replicate 3 (Cook's D = 0.691); normality was restored after its exclusion (W = 0.956, p = 0.135). All VIF values were < 1.1, confirming no problematic collinearity among predictors.

| **Model** | **Shapiro-Wilk W** | **Shapiro-Wilk p** | **Max VIF** | **Cook's D > 0.5** |
| --- | --- | --- | --- | --- |
| Density (acute) | 0.970 | 0.362 | 1.03 | 0 |
| Chlorophyll (acute) | 0.864 | < 0.001 | 1.03 | 1 (C2, r3) |
| Chlorophyll (excl. r3) | 0.956 | 0.135 | — | 0 |

**Figure S2. Residual diagnostic plots for the main stability models.**

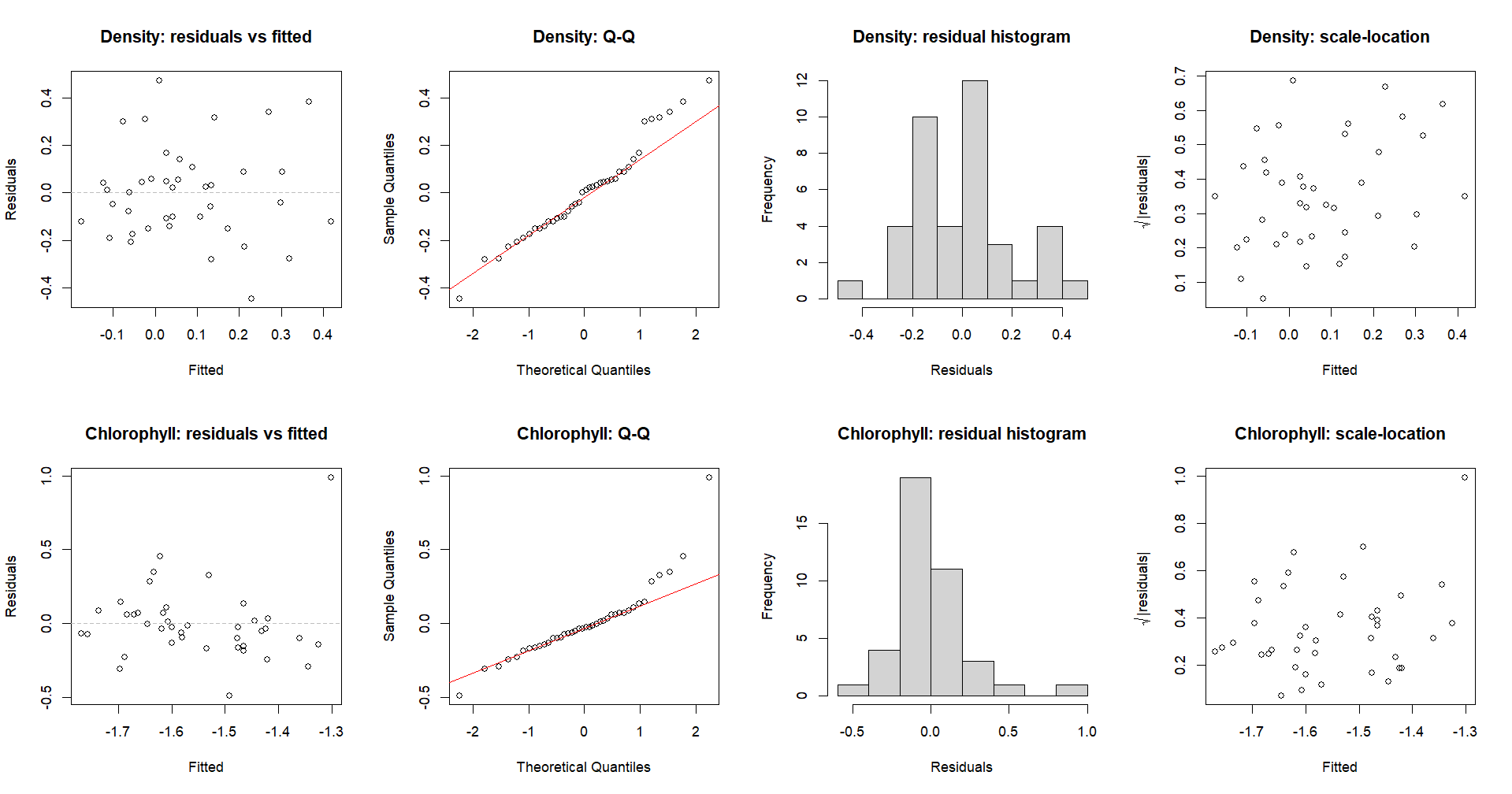

**Figure S2.** Residual diagnostic plots for the log(CV) ~ richness + mean response + response diversity model fitted to total-density stability (top row) and per-cell chlorophyll a stability (bottom row). Left to right: residuals versus fitted values, normal Q-Q plot, residual histogram, and scale-location plot. Density residuals were approximately normal (Shapiro-Wilk W = 0.970, p = 0.362; Table S6). Chlorophyll residuals showed departure from normality (W = 0.864, p < 0.001) attributable to community replicate 3 (Cook's D = 0.691); after its exclusion, residuals were approximately normal (W = 0.956, p = 0.135; Table S6).

**Table 7. Total density stability model: full coefficients.**

Coefficients (β), standard errors (SE), bias-corrected and accelerated (BCa) bootstrap 95% confidence intervals (2000 iterations, community resampling), and p-values for each term in the model log(CV) ~ richness + mean response + response diversity, fitted to total-density stability using acute-derived predictors (main model; R² = 0.36, F₃,₃₆ = 6.61, p = 0.001) and chronic-derived predictors (null comparison). BCa CIs are reported only for the continuous response-metric terms. All predictors were mean-centred prior to modelling. The acute model is the primary result (Fig. 3); the chronic model serves as a pre-specified null showing that chronic metrics do not predict stability.

| **Model** | **Term** | **β** | **SE** | **95% CI (BCa)** | **p** |
| --- | --- | --- | --- | --- | --- |
| Total density (acute) | Intercept | 0.104 | 0.046 | — | 0.031 |
|  | Richness (3 vs 2) | −0.059 | 0.066 | — | 0.375 |
|  | Mean acute response | 0.186 | 0.067 | [0.071, 0.324] | 0.008 |
|  | Acute response diversity | −0.200 | 0.064 | [−0.316, −0.066] | 0.003 |
| Total density (chronic, null) | Intercept | 0.108 | 0.057 | — | 0.068 |
|  | Richness (3 vs 2) | −0.067 | 0.082 | — | 0.419 |
|  | Mean chronic response | 0.047 | 0.115 | [−0.185, 0.280] | 0.685 |
|  | Chronic response diversity | −0.001 | 0.138 | [−0.291, 0.283] | 0.996 |

**Table 8. Chlorophyll a stability model: full coefficients and sensitivity analysis.**

Coefficients (β), standard errors (SE), bias-corrected and accelerated (BCa) bootstrap 95% confidence intervals (2000 iterations), and p-values for each term in the log(CV) ~ richness + mean response + response diversity model fitted to per-cell chlorophyll a stability, using acute-derived predictors (main model), chronic-derived predictors (null comparison), and the acute model with the influential replicate r3 excluded (sensitivity). BCa CIs are reported only for the continuous response-metric terms; indicates CI not computed for intercept or richness. The response-diversity term approaches significance only in the full acute model and is not robust to the influential-point exclusion (see main text).

| **Model** | **Term** | **β** | **SE** | **95% CI (BCa)** | **p** |
| --- | --- | --- | --- | --- | --- |
| Chlorophyll (acute) | Intercept | −1.51 | 0.057 | — | < 0.001 |
|  | Richness (3 vs 2) | −0.086 | 0.081 | — | 0.297 |
|  | Mean acute response | 0.089 | 0.082 | [−0.075, 0.318] | 0.286 |
|  | Acute response diversity | −0.193 | 0.078 | [−0.484, −0.054] | 0.018 |
| Chlorophyll (chronic, null) | Intercept | −1.52 | 0.061 | — | < 0.001 |
|  | Richness (3 vs 2) | −0.076 | 0.088 | — | 0.393 |
|  | Mean chronic response | 0.034 | 0.124 | [−0.218, 0.384] | 0.785 |
|  | Chronic response diversity | −0.195 | 0.148 | [−0.784, 0.091] | 0.195 |
| Chlorophyll (acute, excl. r3) | Intercept | −1.56 | 0.042 | — | < 0.001 |
|  | Richness (3 vs 2) | −0.035 | 0.059 | — | 0.564 |
|  | Mean acute response | 0.032 | 0.060 | [−0.075, 0.318] | 0.601 |
|  | Acute response diversity | −0.109 | 0.058 | [−0.484, −0.054] | 0.071 |

**Text 4:** Derivation of the additive (non-interacting) prediction of community growth, showing why the appropriate combination of strain growth rates is the logarithm of the mean exponentiated rate, ln(mean(exp(ρ))), rather than the arithmetic mean of the rates, mean(ρ).

For strain
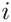
, the per capita growth rate over the interval from
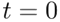
to
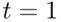
is

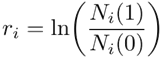

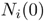
and
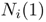
denote the densities at the beginning and end of the experiment, respectively.
To predict the growth of a mixture of two strains in the absence of interactions, we assume that each strain grows independently at its monoculture growth rate and that both strains are initially seeded at the same density

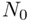
. The expected densities at
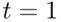
are therefore

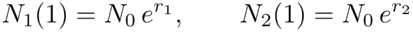

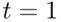
is

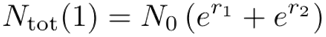

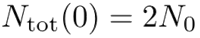

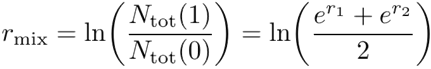

, which is only equal to
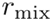
when
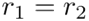
. More generally, the expected growth rate of a non-interacting mixture is determined by the arithmetic mean of the growth factors
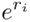
rather than by the arithmetic mean of the log-transformed growth rates.

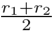

**Table 9. Departure from the additive prediction of community growth, per transition.**

For each of the three thermal transitions in the fluctuation regime, the mean signed departure of observed community per-capita growth rate (ρ_o) from the additive prediction (ρ_p = ln((1/S) Σ exp(ρ_i)), the growth rate of a non-interacting equal-density mixture of constituent monocultures), tested against zero with a linear mixed model (random intercept for community and replicate). Negative values indicate sub-additivity (communities grew more slowly than predicted). Also shown are the slope, intercept, and R² of the within-transition regression of ρ_o on ρ_p, and the number of communities (n) per transition.

| **Transition** | **Mean departure** | **p (vs zero)** | **Intercept** | **Slope** | **R²** | **n** |
| --- | --- | --- | --- | --- | --- | --- |
| C→T (first warming) | −1.26 | 0.001 | −0.509 | −0.289 | 0.028 | 40 |
| T→C (cooling) | −0.17 | 0.660 | 0.812 | 0.101 | 0.009 | 40 |
| C→T (second warming) | −1.12 | < 0.001 | −1.150 | 0.160 | 0.028 | 40 |
